# Decade-Long Impact of Transcranial Direct Current Stimulation in Alzheimer’s Disease

**DOI:** 10.1101/2025.01.26.634937

**Authors:** Anu Aggarwal

## Abstract

Alzheimer’s disease (AD) is a protracted illness with no known cure. Transcranial direct current stimulation (tDCS) is an upcoming treatment for the disease. However, clinical studies conducted to study its impact on patients are very limited in duration and sample size. This limits the use of tDCS to AD patients over the long-term. To address the issue, this work uses an Alzheimer’s disease progression model (ADPM) to extrapolate the results from shorter clinical studies (10days to 6months) to a longer duration (10years). ADPM is a machine learning model of AD created using ADNI (Alzheimer’s Disease Neuroimaging Initiative) data from over a decade. It has been demonstrated to be better at predicting progression of AD compared to other models. Since AD is a progressive disease, it is useful to know about the impact of tDCS over a longer duration and predict the optimal duration and site of treatment. To achieve this goal, present work integrates limited data from tDCS treatment of AD patients and extrapolates it to a decade using the AD prediction model (ADPM). Based on this, it provides guidelines for site, dose and duration of treatment for patients with AD.

## 1. Introduction

Alzheimer’s disease (AD) is the most common cause of dementia, accounting for 60-80% of cases in the US [1]. It is a progressive neurodegenerative disorder that primarily affects memory and adversely impacts the activities of daily living [2]. Despite ongoing research, there is no definitive cure for the disease, leading to high end-of-life health care costs.

According to [3]-[7], AD is characterized by accumulation of extracellular beta-amyloid plaques and neuronal p-tau [8], [9] in the brain. E4 allele of the apoE gene is highly enriched in neurodegeneration associated neuroglia [10]-[12], and increases the protein kinases responsible for tau phosphorylation [13]-[16]. As the disease advances, there is a reduction in brain volume [17]-[24] due to fibrosis after the abnormal protein deposits trigger inflammation. Even though decline in brain volume is observed in healthy controls with ageing (0.5% per year), it is significantly faster in patients with AD (1-3% per year) and the severity depends on whether the AD is familial (early onset, faster decline) or sporadic. Disease diagnosis and progression are indicated by changes in brain size on MRI [25][26], neuronal activity on fMRI [27][28], amyloid and p-tau deposits on PET [29]-[32], and cognitive changes on clinical examination (MMSE) [33]-[35], [4]. These can be used to monitor disease progression alongside things like CSF or plasma levels of amyloid beta 42/40 ratio, p-tau, neurofilament light chains, and glial fibrillary acidic protein [36]-[47]. The disease is also associated with increased metabolic injury and metabolic markers like cytokines, interleukins etc. [48][49]. Various treatments [50]-[54],[4] have tried to target receptors for metabolic markers. Other treatment modalities include gene editing to increase TRIM11 and p-tau clearance, decrease APOE and TREM-2 to reduce beta amyloid formation among others. More recent treatment modalities include deep brain stimulation, transcranial direct current stimulation (tDCS), transcranial alternating current stimulation (tACS), cognitive stimulation, rehabilitation, physical exercise and lifestyle modification [55]-[90],[17]. However, documented use of tDCS treatment [91]-[107] is limited in duration and number of patients due to which it is difficult to assess its impact over a longer duration. Over the short duration of these studies, a significant increase in cognitive scores has been observed in treatment groups as compared to sham, as determined by the Mini Mental State Exam (MMSE) scores. Since AD is a progressive disease, it will be useful to know the impact of any treatment over a longer duration. Work presented here extrapolates the cognitive score data from the short-term tDCS studies to a period of a decade using the Alzheimer’s Disease Progression Model (ADPM)[108].

## 2. Background

The ADPM uses data from the Alzheimer’s Disease Neuroimaging Initiative (ADNI) [109] for 160 patients over a 10-year period. This data contains PET and MRI images and MMSE score of the patient over a decade. The model represents brain as a graph, *G*_*s*_*=(V, E)*, where a node represents either the hippocampus (HC) or the pre-frontal cortex (PFC) and an edge represents the white matter tracts connecting them. *D*_*v*_ (*t*) is the instantaneous amyloid accumulation (determined from PET images) in the brain region, *v* at time, *t*. As the disease progresses, the amyloid propagates to other areas of the brain [110] as

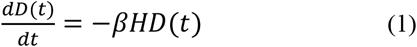

Where *β*is a constant and H is a matrix predicting amyloid propagation between the regions. Amyloid accumulation changes size, *X*_*v*_*(t)* (baseline is taken from MRI images)– initially, increasing it and eventually reducing it [111]. *X*_*v*_*(t)* can predict activity, *Y*_*v*_*(t)* and information processed, *I*_*v*_*(t)* [112], [113]:

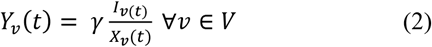

where *γ* is a constant. *I*_*v*_ and *Y*_*v*_ across the 2 regions yield the total cognitive score, *C* and the total metabolic cost, *M*, respectively. *X*_*v*,_ *D*_*v*_ and *Y*_*v*_ are related [114]

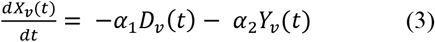

where *α*_1_ and *α*_2_ are constants. The model parameters - α_1_, α_2_, β, γ were calculated from the demographic features of the population in ADNI. Since *I* could not be derived from ADNI data, reinforcement learning (RL) was used to find it by maximizing the following reward, *R*.

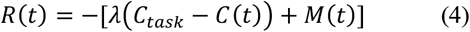

where *C*_*task*_ is the cognitive demand of a task. *Y* depends on *C*_*task*_ [115] and has an *M* which can enhance degeneration [114]. In the RL, the state consists of *X(t)* and *I(t-1)*. The environment is a simulator comprising the above equations. Action, *ΔI(t)* is the change in *I(t)*. Based on the action, the environment updates the state and provides a reward. The RL agent was trained on the simulator with model free on-policy learning using TRPO for 1 million epochs of batch size 1000. There were 32 hidden units in each of the two-layer feedforward network implemented using Open AI’s Gym framework. λ and *I*_*o*_ were chosen using grid search to minimize the validation set error, *C* was normalized between 0-10, *C*_*task*_ was fixed at 10 and *X* was normalized to 3.5-5.5. ADPM successfully modelled disease progression over a 10-year period with lower error than previous models like the miniRNN [116], support vector regression [117] and [110]. We used the ADPM model to integrate the findings from tDCS clinical trials and run it to provide disease progression with tDCS.

The MMSE [118] is a clinical exam in which patients are administered 11-questions to evaluate their cognitive functions-orientation, registration, attention, recall and language. The score varies from 0 to 30, 30 being the best.

In tDCS, a constant (direct current or DC) electrical current (∼2mA) is passed through anodal electrodes placed on scalp (like EEG electrodes) such that a specific brain area can be stimulated [119]. The tDCS could have local impact on the GABA/glutamate balance - anodal tDCS decreasing and cathodal increasing the GABA-ergic inhibition [120] to influence functional connectivity, synchronization, and oscillatory activities in prefrontal cortex in healthy individuals as determined by EEG and low resolution tomography [90], cause glial stimulation [121] which leads to neurotransmitter release, or could modulate the conformation of beta-amyloid proteins [122]or increase LTP-like plasticity [123] in human cerebral cortex. Priors from [91]-[107] reported change in MMSE scores of AD patients treated with tDCS over a short duration (Table 1).

**Table 1.** Summary of all the priors used in integrating the change in MMSE scores in this study.

| Study | Treatment group (number) | Sham group (number) | Duration | Region stimulated | Change in MMSE score after days of treatment |  |
| --- | --- | --- | --- | --- | --- | --- |
|  |  |  |  |  | Treatment group | sham group |
| Khedr 2019 | 23 | 23 | 10 sessions of 2mA, 20mins a day, 5 sessions a week, for 2 weeks | Left temporoparietal region | 10d=+3.5 | 10d=-0.6 |
| Khedr 2014 | 11-atDCS<br>12-ctDCS | 11 | for 25 min at 2 mA daily for 10 days | left dorsolateral prefrontal cortex | 10d = +1.6, 1mo = +4, 2mo =+4 | 10d=1.5, 1mo= -0.5, 2mo=-1 |
| Gange mi et. al. | 13<br>9 | 13<br>9 | Anodal tDCS 2mA for 20minutes a day for 10d<br>10 days a month for 8m. | left frontotemporal lobe | -0.3<br>-0.28 | -2.4<br>-4.08 |
| Cotelli et. al. | tDCS+memory training=12<br>tDCS+motor training=12 | 12 | 10 sessions of 2mA tDCS for 25mins a day, 5 sessions a week, over 2 weeks | Dorso lateral pre-frontal cortex (DLPFC) | tDCS+memory 2wks=+0.5, 3mo=-0.4, 6mo=-0.5<br>tDCS+motor 2wks=+0.2, 3mo=-1.1, 6mo=-0.3 | 2wks=+0.9, 3mo=-0.3, 6mo=+0.2 |
| Gronli et. al. | 8 | Self-control | 2mA of anodal tDCS was applied for 30minutes daily for 4months | right DLPFC | +1, -1 after 4months and 8months of study start | NA |
| Im et. al | 11 | 7 | every day for 6 months (anode F3/cathode F4, 2mA for 30 minutes) | DLPFC | +1.1 | -1.5 |
| Boggio et. al. | 15 | 15 (self-control) after 71-77 days | 2 mA, 30 min per day for 5 days | bilaterally over the temporal regions | +0.1, 0.9, -0.1 at 0, 1, 4weeks after treatment | +0.3, 0.7, 0.2 at 0, 1, 4weeks after treatment |
| Bystad, et. al. | 12 | 13 | 2mA in six 30-minute sessions over 10 days. | left temporal cortex | +1 | +1 |
| Gomes et. al. | 29 | 29 | 2.0 mA, for 30 mins for 10 sessions, twice a week | left DLPFC | +0.3 | No change |
| Chu et. al. | Meta analysis of 27 RCTs |  |  |  | <1mon=+0.56, >1mon=+0.21 | NA |
| Lu et. al. | 61 in tDCS and working memory training (WMT)group, 54 in tDCS and control cognitive training (CCT) group | 58 in sham tDCS and WMT | for 20 min at 2 mA, with three 45min sessions per week, for 4 weeks | left lateral temporal cortex | After 4, 8 and 12 wks- 1.14, 0.57, 0.32 (with WMT)<br>0.68, 0.52, 0.71 (with CCT) | 0.66, 0.65, 0.14 after 4, 8 and 12 wks |
| Fileccia et. al. | 17 | 17 | 2-mA for 20 min, 5 days per week (total=20 days) | left DLPFC | +0.7 | -0.5 |
| Hu et. al. | 21 in rTMS-tDCS<br>21 in rtMS<br>21 in tDCS | 21 | 30 minutes stimulation on both angular gyri for 4 weeks | Angular Gyrus and prefrontal area | 4.86, 2.62 for rTMS-tDCS<br>3.62, 1.62 for rTMS<br>3.28, 1.38 for tDCS at 4 and 12 weeks | +0.2, -1.1 at 4 and 12 weeks |
| Inagawa et. al. | 10 | 9 | 2mA, 20 min, twice daily for 5 consecutive days | DLPFC | 0.41, 1.08 more than sham at day 0 and 2 weeks after | NA |

Above we summarize how the ADPM provides AD projection over a decade supported by ADNI data set, the limited clinical studies on tDCS and indication of clear benefit of tDCS in AD from them. Thus, to understand the long-term impact of tDCS in AD, we modified the ADPM in the current work, to incorporate the impact of tDCS stimulation of either the HC, the PFC or both over a period of a decade.

## 3. Methods

The ADPM processed ADNI data using domain knowledge guided differential equations and RL to predict change in cognitive scores for AD patients over a decade. Thus, to integrate the impact of tDCS, we had to incorporate the parameters that tDCS studies use into the model parameters. Therefore, there were design decisions to be made.

### 3.1 Design decisions for the intervention

Inputs from the ADNI data to ADPM were:

1. Parameters for the differential equations like *α1, α2, β, γ* and *H* were calculated from patient demographics like age, gender, genetic markers etc. Thus, the model captured the demographic characteristics of the population from ADNI.
2. *D(t)*, determined from PET images at baseline
3. *X(t)* from MRI data at baseline
4. *C(t)* from MMSE scores at baseline

Based on data inputs, domain knowledge guided equations and RL agent, the model predicted change in *X(t)* and *I(t)* at one-year intervals over a decade and compared it to the ADNI data. *I(t)* was contribution of a region to the cognitive score. The priors [91]-[107] reported the dose and duration of tDCS and the impact of treatment on MMSE scores. The ADPM does not use the tDCS current as input to the model. The only input to the model that changed yearly was *I(t)* obtained from RL. This was integrated into the model at each timestep and the corresponding change in brain volume (*X(t)*) was reported at each time step. The *X(t)* has been demonstrated [111]-[113] to be linked to disease progression. Thus, to integrate findings from priors into the model, we integrated the impact of tDCS as change in MMSE scores reported in the studies as new *ΔI*_*stim*_ at each time step. And we observed the change in *X(t)* to determine the impact on disease progression-increase in *X(t)* meant that the disease progression has reversed. The details of the intervention are provided in the next section.

### 3.2 Modification Applied to the Model

We modified the ADPM to introduce the effect of tDCS on a patient as improvement in cognitive score for a given current applied based on data from priors (Fig. 1). For this, a new stimulation parameter, Δ*I*_*stim*_ was added to the model. In this updated version of the ADPM, a new version of *I*_*v*_(*t*) called *Îv*(t) is defined as:

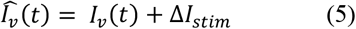

*Î*_*v*_(*t*) replaces *I*_*v*_(t) in equation (2) in the ADPM. Since *Y*_*v*_(*t*) will change due to this addition, the impact will propagate to calculation of X_*v*_(*t*) in (3). An increase in brain volume indicates improvement in brain function and a decrease in volume indicates reduction in brain function. We were interested in determining

- What region should be stimulated with 2mA tDCS for 30minutes a day for 10days a year for best effects?
- How much of tDCS should be administered?

1. To determine the region to be stimulated for optimal effect, we used a Δ*I*_*stim*_ value of 0.8, which equates to an MMSE score increase of 2.4 per year. This level of Δ*I*_*stim*_ was chosen because the stimulation data from ***Table 1*** indicates an average increase of 2.4 points on the MMSE per year with 2mA for 30 minutes a day for 10days. The change in MMSE score was integrated into the model using the method described above. Upon stimulation, change in brain region volume (HC or PFC) over a period of 1-10 years was studied as an indicator of disease progression. Six stimulation regimens were used:

- 3years of HC stimulation followed by 3years of PFC, followed by 4years of HC (HC334),
- 3years of PFC stimulation followed by 3years of HC, followed by 4years of PFC (PFC334),
- 5years of PFC followed by 5years of HC (PFC5),
- 5years of HC followed by 5years of PFC (HC5),
- 2years of alternating HC and PFC stimulation – beginning with
  - HC (HC2)
  - PFC (PFC2)

2 To determine how much tDCS is required, a range of Δ*I*_*stim*_ between 0.1 to 5 was applied to the model. This corresponds to a range of MMSE score increase between 0.3-15 (as the cognitive score is normalized to 0-10 in the model instead of 0-30).
3 To determine the region of brain that needs stimulation, it was applied to either the PFC or the HC or both. When applying Δ*I*_*stim*_ to both regions, the value of Δ*I*_*stim*_ applied to each region was halved.

**Fig. 1.**
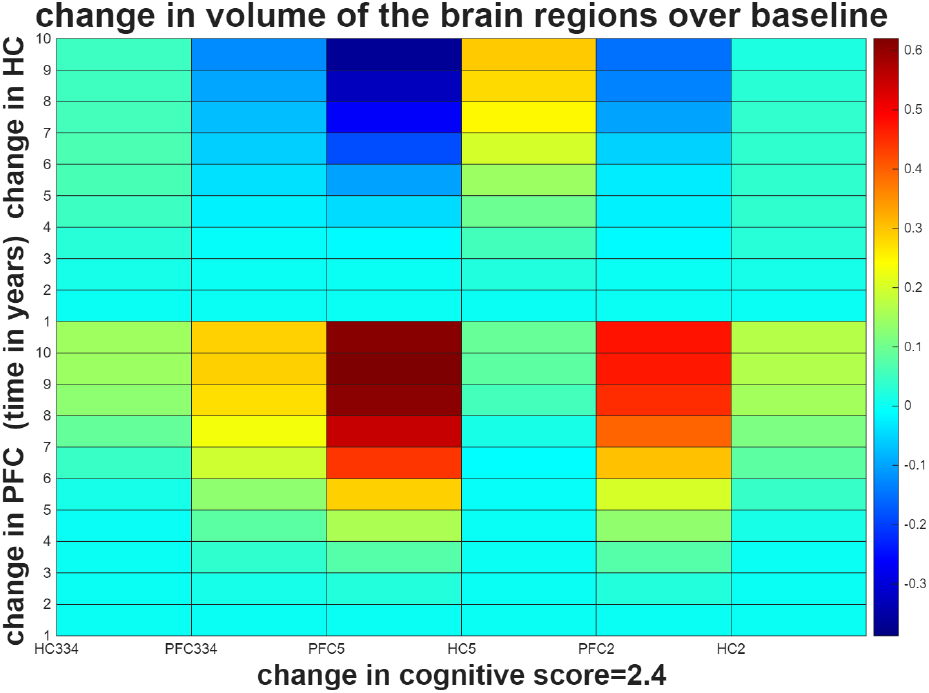
Change in mean PFC and HC volumes when alternating PFC and HC stimulation over time. Standard deviations are represented as error bars. Differences are with respect to baseline results at the same year for untreated patients.

As noted in [112][113], brain structure predicts brain activity which results in cognition which is measured using tests like the MMSE. Thus, in the present work, we use change in brain size to determine the resulting MMSE score after application of intervention (2mA of tDCS for 30 mins daily). The results from different stimulation paradigms are shown in the next section.

## 4. Results

The original ADPM provides projected values for brain volume over 10 years, serving as a baseline to quantify an increase or decrease of region volume. The mean values of the difference in volume between baseline and stimulation results are shown in Figs 1-3.

### 4.1 Alternating PFC and HC Stimulation Over Time

For this experiment, the Δ*I*_*stim*_ per year was 0.8, as average of priors results in an MMSE increase of 2.4 per year. Six different stimulation protocols were used with this Δ*I*_*stim*_ as mentioned earlier. From Fig. 1, it is evident that the maximum change in non-normalized PFC volume at year 10 was 16755 mm^3^, which is like that of 17502 mm^3^ with PFC stimulation alone (shown in Fig. 2). This maximum was achieved in the PFC5 protocol. The dual stimulation (as seen in Fig. 3) for 2.4 projected increase in MMSE score resulted in a maximum PFC volume increase of 10,826 mm^3^. This indicates that to get the maximum improvement in health of the PFC, the tDCS should be applied to the PFC alone. From the results, it is also evident that for the HC, the maximum unnormalized volume change was 594 mm^3^ using the HC5 protocol. For the 2.4 projected MMSE score increase, the maximum change is lower than that from single stimulation (1013 mm^3^), but higher than that from dual stimulation (-108 mm^3^). Therefore, to attain the maximum HC volume increase, stimulating only the HC seems to be the best option. From the figure, it is also evident that the stimulation protocol that results in the largest increase in PFC volume results in the largest decrease in HC volume. The protocol with the largest increase in HC volume does not decrease, but it attains the smallest increase in PFC volume by year 10. Thus, making it difficult to attain best results for both regions with a single stimulation protocol.

**Fig. 2.**
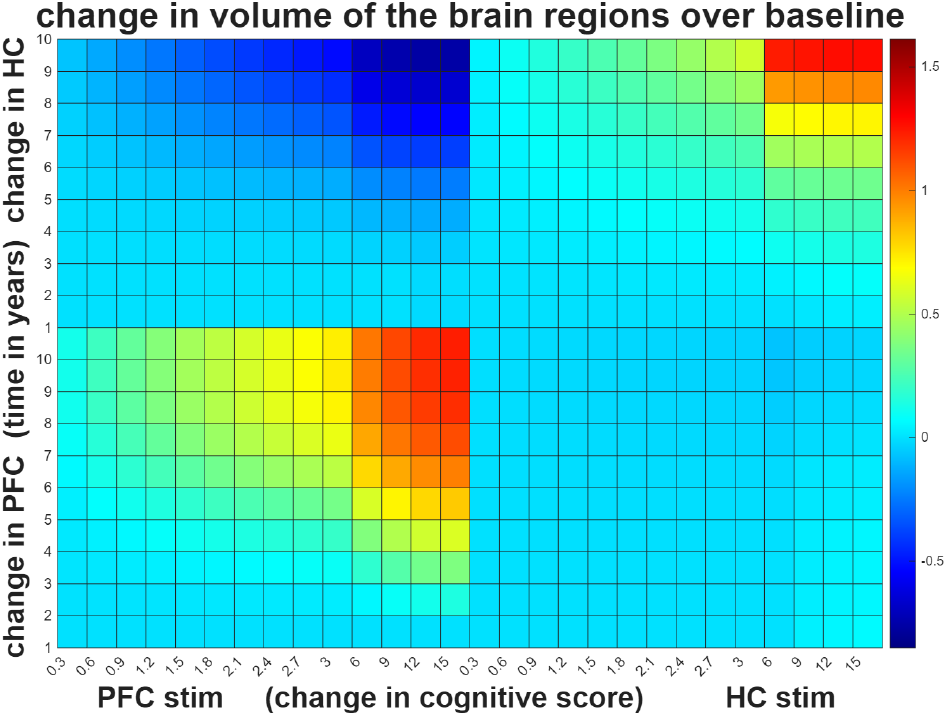
Showing mean change in HC and PFC volume over a period of 10 years when different current stimulation and corresponding change in MMSE scores was applied over a period of 10years in the ADPM. Change in volume is with respect to baseline results for the same year when patient did not receive any treatment. There is a positive volume change in the PFC or the HC if the stimulation is applied to the same region as opposed to that applied to the other region. HC stimulation does not seem to have a negative impact on the PFC volume, but PFC stimulation has a negative impact on the HC volume.

**Fig. 3.**
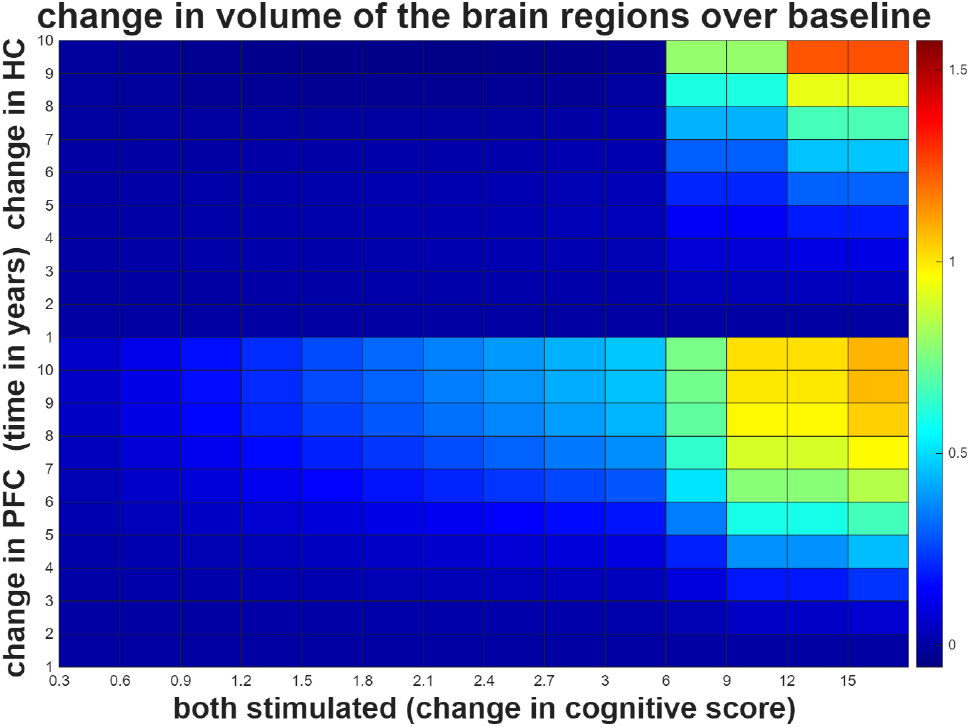
Mean change in PFC and HC volume over a period of 10 years when different current stimulation and corresponding change in MMSE scores was applied in the ADPM – with half the stimulation applied to HC and the other half to the PFC. Differences are with respect to baseline results at the same year for untreated patients. There is a positive volume change in both the PFC and the HC if the stimulation is applied to both regions but only in the later years and with much higher level than when stimulation is applied to either region alone.

### 4.2 PFC and HC Stimulation

The largest changes in PFC or HC volume occur when stimulating only the respective region. Across all points and levels of stimulation, the PFC volumes increase over baseline when stimulating the PFC, and the HC volumes increase over baseline when stimulating the HC. Furthermore, the value of Δ*I*_*stim*_ largely impacts the rate of volume change. Stimulating the PFC tends to produce a decrease in HC volumes over time, as shown in Fig. 2. And when HC stimulation is applied, there is a noticeable reduction in PFC volume relative to baseline across all MMSE score improvements (in the range of 124 mm^3^ to 1887 mm^3^, with higher MMSE score improvements leading to larger decreases). There is improvement in brain volume for the region with as little as 0.6 and 0.9 increase in Δ*I*_*stim*_ for PFC and HC respectively. For smaller values of Δ*I*_*stim*_, the impact is visible later indicating a cumulative effect over time. This indicates we could use a smaller or larger dose of tDCS and get the impact on MMSE scores at a different time. This could also be modulated based on the initial MMSE scores and needs of the patient.

### 4.3 Simultaneous PFC and HC Stimulation

In this experiment, for each value of Δ*I*_*stim*_, half of the stimulation was applied to the HC and half to the PFC. The overall result trends are like those in the second experiment, but differ in magnitude, as seen in Fig. 3. Dual stimulation produces a dampened version of the results of single region stimulation. For example, with a projected cognitive score increase of 0.6 per year, the dual stimulation results in a mean PFC volume increase of 3173.14mm^3^ over baseline at year 10, whereas stimulating only the PFC with the same projected cognitive score increase results in an increase of 6054.72 mm^3^ in actual (not normalized) volume. All projected cognitive score increases result in an increase in the PFC volume over baseline at the same time for all timesteps. Lower projected cognitive score increases per year result in decreased HC volumes over baseline. By year 10, MMSE score increases of 6 per year (corresponding to Δ*I*_*stim*_ of 2.0) or higher are the only ones causing an increase in the HC volume over baseline. But one notable difference from simulations in section 4.1 and 4.2 is that there is no decrease in cognitive scores or region volumes in any of the regions – HC or PFC when both are stimulated at the same time.

## 5. Discussion and Conclusion

The inclusion of a stimulation parameter in the ADPM models the effect of tDCS on AD patients. The model determines how different cognitive score increases per year can affect brain region volumes over a decade. Based on the results, the single region stimulation seems to be the most effective at increasing volume of the region among all the stimulation protocols while dual stimulation also increases the volume but at a slower rate. However, dual stimulation has the advantage of not reducing any region’s volume. Alternating stimulation can produce comparable results in the case of PFC volumes when compared to single region stimulation, but at the expense of decreasing HC volumes. Given that higher brain volumes are linked to increased brain activity and better cognitive health [107], dual stimulation simultaneously seems to be better over alternatives. Our simulations indicate that lower tDCS dose will also improve brain region volume and hence, cognition like higher doses used in priors but over a longer duration. For patients with mild AD, perhaps smaller doses of tDCS will be useful. Our simulations also indicate cumulative effect of treatment over time. Therefore, we recommend clinical trials with smaller doses, dual region application and longer duration of tDCS to validate the results from this model-based simulation.

To conclude, this work examines the impact of tDCS stimulation on brain volume and severity of disease in AD patients. It utilizes an existing model to simulate AD progression with tDCS over ten years by modifying the model based on priors. This work also demonstrates a novel modality, viz., building model of a disease using patient data and machine learning algorithms to test treatments which can’t be tested clinically or as a prior to large-scale clinical trials.

